# Genetically mediated associations between chronotype and neuroimaging phenotypes in the UK Biobank: a Mendelian randomisation study

**DOI:** 10.1101/2023.08.31.555801

**Authors:** John A. Williams, Dominic Russ, Laura Bravo-Merodio, Georgios Gkoutos, Mark A. Bellgrove, Andrew P. Bagshaw, Magdalena Chechlacz

## Abstract

Chronotype impacts numerous physiological and disease traits, from metabolic syndrome to schizophrenia. The suprachiasmatic nucleus (SCN) maintains transcriptional-translational feedback loop (TTFL) which acts as a central chronobiological pacemaker, regulating 24-hour cycles throughout the human body. However, each tissue maintains its own peripheral clock, and both endogenous hormones and neurotransmitters and exogenous environmental cues regulate the SCN’s central clock. The extent to which brain regions outside the SCN influence the core TTFL is unknown. Here, we investigated how genetic variability affecting brain regions outside the SCN may indirectly influence chronotype, using Mendelian randomization and causal inference. We performed genome wide association studies (GWAS) based on image derived phenotypes (IDPs) from neuroimaging data (grey matter volume, thickness and surface area, microstructural white matter measures; 42,062 participants), and additionally for sleep duration and morning/evening chronotype (361,739 participants). Significant, single nucleotide polymorphisms (SNPs) associating with each phenotype were entered into 2-sample Mendelian randomization performed using inverse-variance weighted methods (exposure versus outcome): 1) chronotype versus each IDP, 2) sleep versus each IDP and 3) each IDP versus chronotype. Subsequently, we investigated genes where significant instrumental SNPs were located for circadian periodic cycling, interaction with TTFL genes in common biological pathways (genetic, physical, or functional interaction), and enrichment of traits from UK Biobank and GWAS Catalogs. We found three associations with chronotype (morning/evening diurnal preference) outside the SCN based on genetically predicted (FAM76B, DENND1A, CDH11) regional differences in brain volume. Specifically, genetically predicted lower inferior temporal gyrus volume linked to morning phenotype, while lower volume of the superior parietal lobule and angular gyrus linked to evening preference. In addition, evening chronotype exposure influenced superior temporal gyrus volume, and both increased sleep duration and evening chronotype influenced thalamic volume. We conclude that genetically mediated associations between chronotype and brain regions outside SCN exist suggesting novel zeitgeber mechanisms.

## Introduction

Nearly every cell in every organism on the planet has a circadian rhythm, a biological process which oscillates over a 24-hour period. These endogenous circadian rhythms are internally self-regulated, occurring in the absence of any external influences, acting in anticipation of daily environmental changes to which they are highly susceptible (for recent review see [1]). In mammals, the phase of circadian rhythm can be set by external cues called zeitgebers, or”time givers” [2]. The zeitgebers are predominantly exogenous environmental cues acting as repetitive oscillators, with daily changes in light being the most powerful zeitgeber [3]. Light acting as zeitgeber, enters the retina and entrains the core mammalian clock in the suprachiasmatic nucleus via the optic nerve and the circadian entrainment molecular pathway. The core molecular clock consists of transcriptional-translational feedback loop (TTFL) with translated clock proteins (CLOCK, BMAL1) acting as transcription factors for additional clock genes (*Per1/2/3,Cry1/2*). These genes, once translated and dimerized, negatively inhibit the action of CLOCK and BMAL1 before being phosphorylated and degraded. The entire process takes roughly twenty-four hours [4]. This central clock acts as a”pacemaker” of tissue-specific, peripheral clocks throughout the rest of the body [**?**, 5]. The most widely studied peripheral circadian clock is in the liver, where metabolism is affected by clock genes [6–9]. There is also increasing evidence that peripheral clocks exist within mammalian brain [10], with numerous core clock genes known to be expressed in the rodent cerebral cortex and hippocampus [**?**, 11, 12]. While further work is needed to fully understand the function of these peripheral clocks, the current understanding is that they likely act as internal cues, exerting influence on body temperature or over the levels of hormones, neurotransmitters, or metabolites [12, 13], and thus affecting body biochemistry, physiology, and behaviour.

The circadian cycle regulated by SCN and TTFL always oscillates within approximate 24h period, however a large variability exists in the actual timing of the clock generated biological processes, including sleep-wake cycles. This variability is commonly referred to as chronotype, a diurnal morning versus evening preference in sleep-wake phases and other associated biological processes. While many core circadian clock genes are well known, the genetic underpinnings of diurnal phenotypes remain to be fully understood and catalogued, with several recent GWAS studies providing essential first insights into genetic variance underlying chronotype [14, 15]. Recent GWAS of self-reported chronotype (lark or owl, corresponding to being a ‘morning person’ or ‘night person’) revealed several candidate genes associations with previously unexpected circadian function [16].

Genome-wide association and mutation studies have linked variants in core clock genes to sleep-wake disorders, disruptions due to jet lag, and even sleep-related bone loss [17]. Abnormalities of circadian rhythms and variation in diurnal preference are associated with a myriad of neurobehavioral disorders in human, from schizophrenia to major depressive disorder [18–21]. The translational importance of understanding biology of diurnal preference is not limited to neurobehavioral function; the interplay between metabolism and chronotype has been highlighted heavily in recent years. The gut microbiome, adipose cytokines, and metabolic hormones from ghrelin to leptin are all strongly regulated by circadian biology [22–24] as well as linked to chronotype [25, 26]. Strikingly, the existing literature supports a notion that a bidirectional interplay between gut hormones regulating metabolism and chronotype influence neuropsychiatric phenotypes [26].

The extent to which brain regions outside the SCN influence the core TFFL and diurnal preference is unknown. However, with the known links between chronotype and multiple behavioural and psychiatric traits, it is plausible that a relationship between diurnal preference and inter-individual regional brain variance exists, which might in turn mediate some of the observations (i.e., links between diurnal preference and psychiatric phenotypes).

Recently a large-scale epidemiological study, UK Biobank, with extensive behavioural questionnaire, neuroimaging, and genetic data has been created [27]. This unique resource facilitates examining multifactorial interactions using statistical genomics approaches on a scale not feasible before. Recent studies using early UK Biobank data release (first 5,000 participant from the UK Biobank Brain Imaging Cohort [28], demonstrated link between self-reported diurnal preference and volume of numerous brain regions, including hippocampus, orbitofrontal cortex and several subcortical nuclei (nucleus accumbens, caudate, putamen and thalamus), subsequently linking some of these volumetric differences to loneliness and depression [29–31]. The current study aims expand these findings and provide novel insights into genetically mediated associations between chronotype and brain regions outside SCN. Thus, we investigated how genetic variability affecting brain regions outside the SCN may indirectly influence chronotype, using Mendelian randomization and causal inference.

Hypotheses: Regional Brain structural differences outside the SCN may influence chronotype Rationale: By investigating genetic-based influences of brain region and diurnal preference via MR, we suggest potential directionality of effect – intracranial (non-scn) zeitgebers. MR robust to statistical signals of heterogenetity or horizontal pleiotropy may suggest chronotype influences indepentent of environmental zeitgebers — activity of central, but non-scn, clocks/feedback.

## Materials and Methods

### 0.1 Data and Participants

Data were taken from the UK Biobank, population-based epidemiological study [27] gathering data on around 500,000 United Kingdom (UK) residents who were aged between 40 and 69 years at study baseline recruitment between March 2006 and December 2010 from 22 centres in the United Kingdom (England, Scotland and Wales). This study utilised UK Biobank collected genetic, brain imaging, sleep and demographic data as systematically described below. Blood sampled collected at baseline were genotyped using two different arrays, the UK BiLEVE Axiom array and the UK Biobank Axiom array (see below). On average 4 years after the baseline recruitment a subset of participants (age range 44-79 years) completed magnetic resonance imaging (MRI) assessment. Genome Wide Association Studies (GWAS) on brain imaging phenotypes (see below), and additionally on sleep and chronotype measures were performed in two non-overlapping participants samples (n = 42,062 [brain imaging cohort], n = 361,739, respectively). The detailed description of UK Biobank recruitment, ethical approval and data has previously been published [27]. All UK Biobank participants provided written informal consent in accordance with approved ethics protocols (REC reference number 11/NW/0382). The described here analyses were conducted under the UK Biobank application numbers 29447 (approved on 2018/09/03) and 31224 (approved on 2018/02/02).

### 0.2 Chronotype Assessment

To collect self-reported measure of diurnal preference (chronotype) UK Biobank participants were asked a question”Do you consider yourself to be?” and asked to choose one of six possible answers:”Definitely a ‘morning’ person”,”More a ‘morning’ than ‘evening’ person”,”More an ‘evening’ than a ‘morning’ person”,”Definitely an ‘evening’ person”,”Do not know” or”Prefer not to answer” (data-field 1180). After exclusion of missing data and participants who answered”Do not know” or”Prefer not to answer” coded as missing, for the purpose of GWAS we generated binary (morning/evening) phenotypes with participants answering”Definitely an ‘evening’ person” and”More an ‘evening’ than a ‘morning’ person” coded as evening type and those answering”Definitely a ‘morning’ person” and”More a ‘morning’ than ‘evening’ person” coded as morning type. For the purpose of GWAS, we also included a measure of sleep duration (data-field 1160). To assess sleep duration participants were asked a question”About how many hours sleep do you get in every 24 hours? (please include naps)”. Participants were asked to input a number using a touchscreen or alternatively provide one of the two answers”Do not know” or”Prefer not to answer”. In addition, the following checks were performed to ensure accuracy: (1) if given answer *<* 1 then rejected, (2) if given answer *>* 23 then rejected, (3) if given answer *<* 3 then participant asked to confirm, (4) if given answer *>* 12 then participant asked to confirm.

Participants were also provided with a”help button”, which showed a message”If the time you spend sleeping varies a lot, give the average time for a 24 hour day in the last 4 weeks”.

### 0.3 MRI data acquisition and Imaging-derived Phenotypes

The neuroimaging data were acquired between 2014 and 2020 (the UK Biobank Brain Imaging Cohort) initially on a single scanner in Cheadle Manchester, and from 2017 in two other identical centres (in Newcastle and Reading). All brain imaging data were acquired on 3T Siemens Skyra Scanners equipped with a standard 32-channel RF receive head coil. Available online UK Biobank Brain Imaging documentation (https://biobank.ndph.ox.ac.uk/ukb/ukb/docs/brain_mri.pdf), and previously published primary UK Biobank methods papers [28, 32] provide full information about the MRI data acquisition protocols and data processing pipelines. The MRI data processing pipeline include robust quality control and processing steps automatically generating modality specific imaging derived phenotypes (IDPs). For the purpose of GWAS we used several structural (grey and white matter specific) IDPs (see Supplementary Table 1 for the full set of employed here IDPs) derived from T1-weighted (a three-dimensional rapid gradient echo sequence with 1mm isotropic voxel resolution acquired in a sagittal plane), and diffusion MRI (a spin echo multiband echo-planar sequence with 2mm isotropic voxel resolution and 100 unique diffusion encoding directions, 50*xb* = 1000*s/mm*2*and*50*xb* = 2000*s/mm*2) scans. Grey matter specific IDPs representing estimates of regional (fronto-parieto-temporal) cortical volume, area and thickness, and thalamic nuclei volumes were derived using various FSL (https://fsl.fmrib.ox.ac.uk/fsl/fslwiki/) and FreeSurfer (http://surfer.nmr.mgh.harvard.edu/) tools as previously described [32], all these measures were normalised for head size prior to GWAS. Following automated tractography [33, 34] (*BEDPOSTx/AutoPtx*) white matter IDPs reflecting microstructural properties of fronto-parieto-temporal pathways were derived from the diffusion tensor imaging (DTI) fitting [35] and neurite orientation and dispersion imaging (NODDI) modelling [36]. This protocol generated white matter IDPs representing fractional anisotropy (FA), mean diffusivity (MD), axial (AD) and radial diffusivity (RD) from DTI model as well as intra-cellular volume fraction (ICVF; a measure of neurite packing density), and the orientation dispersion index (ODI; a measure of dispersion of neurites) from NODDI model. In the GWAS we also included total brain, grey matter, white matter, left and right thalamus volume (all normalised for head size), totalling 293 structural IDPs used here as listed in Supplementary Table 1.

### Genome-wide Association Studies

Imputated genotype data were obtained from UK Biobank and previously described [**?**]. For each variant, additional quality control was performed using Plink v1.90b6.21 and in R [**?**]. Variants with missingness *>* 0.01 and individuals with missingness *>* 0.05 were removed. The self-identified”White” ethnic group was retained to reduce ancestry-based heterogeneity. A 10kb window with step size 5, p1 of 0.99 and p2 of 0.001 were used to remove variants in high linkage disequilibrium (LD) before analysis. Variants with a minor allele frequency *<* 0.01 were excluded, as were those with a Hardy Weinberg Equilibrium of *p <* 1*e −*8. To correct for population structure, the first 10 pre-computed Principal Component loadings from UK Biobank were used as covariates, along with self-reported sex, age, *age*^2^, intracranial volume (applicable brain measures only), and imaging centre location (applicable brain measures only). To perform GWAS associations, Regenie v1.0.6.7 was used [37]. Bgen files were analyzed in a 2-step process of whole-genome regression for linear traits (brain measures and sleep duration), and firth logistic regression was used to estimate a dichotimized morning-evening chronotype. As stated above, individuals with brain phenotypes measured were included in brain measurement GWAS, and remaining individuals were kept for a separate GWAS of chronotype and sleep, resulting data suitable for a 2-sample Mendelian Randomization post-GWAS study.

### Mendelian Randomization

Mendelian Randomization (MR) experiments seek to find evidence of causal relationships between the genetic predisposition of an exposure and and an outcome [38]. Here, we sought to discover relationships between genetically mediated volume change in brain regions outside the SCN and preference for an eveningness (vs morningness) self reported chronotype. For each brain region, significant loci were used as exposure SNPs, and the corresponding *β* effect sizes from chronotype were used as outcome SNPs. The ratio between exposure and outcome, given certain assumptions, suggest evidence for causal relationships.

Since no public GWAS had been preformed for each of the fine-grained subcortical regions analyzed, discovery GWAS were used as exposure SNPs against chronotype. SNPs passing an 5e-8 threshold which were biallelic and non-pallendromic were clumped, using the default settings for the TwoSampleMR package, which was used to conduct MR analyses [**?**]. Independent SNPs, as instrumental variables, were then aligned to the UK Biobank chronotype GWAS, and effect sizes standardized to betas for MR analyses. For a secondary MR, chronotype and a related trait, sleep duration, were exposures against brain region phenotypes.

GWAS results for each outcome/exposure pair (chronotype to brain regions; sleep to brain regions; brain regions to chronotype) were analyzed as individual experiments. For each instrumental variable (IV) SNP in the exposure study, the following procedure was performed. First, the strandedness of each GWAS was checked to ensure that at each allele, the minor and major alleles were equal. If these were reversed, effect sizes were modified to correct for this. Pallendromic SNPs, which contain alleles represented by the same base pairs on both strands of DNA, were discarded. SNPs in the exposure GWAS set were clumped by linkage disequilibrium to ensure statistical independence. In a window of 10,000 base pairs, an *R*^2^ cutoff of ¡ 0.001 was set to obtain haplotype blocks using the European reference panel of the 10,000 Genomes Project [39].

Chronotype effect size was transformed into a *β* from the *log*(*OR*). The Wald ratio was then obtained, giving a measure of the effect of the exposure on the outcome [40]:

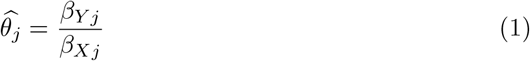

where *β*_*Y j*_ is the effect of the IV on the outcome, and *β*_*Xj*_ is the effect of the IV on the exposure is obtained for SNP j, and 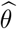 is the effect size.

An initial IVW analysis between each exposure/outcome set was performed. Rather than calculate Wald ratios individually, the outcome GWAS *β*s were regressed on the exposure in an inverse variance weighted (IVW) meta-analysis with both fixed- and random-effects assumptions. The slope of the regression line indicates the strength of the effect, as an increase in the unit of outcome per unit of the exposure [41]. In a IVW meta-analysis, the IVW estimate is calculated by:

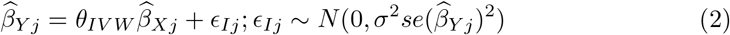

where 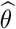 is the inverse variance weighted average, *se* is the standard error, *ϵ* is an error term. To address unseen horizontal pleiotropy, the Egger regression was performed. [42, 43]. The Wald ratios of each SNP are used in meta-regression by taking the inverse variance weights used in the IVW analysis without modeling the intercept. As a result, a causal estimate, similar to IVW, is obtained, adjusted though for horizontal pleiotropy which is necessary so as to ensure IVW validity [43]. MR-Egger regression is an extension of IVW regression. Instead of assuming no intercept term, an intercept is estimated:

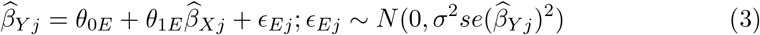

where *θ*_0*E*_ is the intercept and *θ*_1*E*_ the MR-Egger estimate. If the intercept is equal to zero, then the IVW method and MR-Egger will be equivalent [41]. Given the small number of SNPs as potentially valid IVs in some analyses (2-4), we additionally performed a median-based weighted analysis [**?**].

Heterogeneity can occur when individual SNPs do not converge on an estimate; this was estimated by Cochran’s *Q* [44]. In this context, heterogeneity may be a sign of horizontal pleiotropy, wherein SNPs effect the outcome by their influence on other confounding traits [45]. To inspect SNPs for outliers, we performed leave-one-out sensitivity analyses using the IVW method, leaving out one SNP in each analysis (supplement).

As the segmented MRI regions were all non-independent, a strict Bonferroni GWAS would lead to potential loss of power owing to the independence assumption [46]. To threshold the family wise error rate, we performed a principal component analysis on all IDPs where MR analyses was performed. Thresholding to explain 90% variance yielded 46 components, for a p-value threshold of 0.05*/*46 = 0.001. Any MR with an IVW test ¡ 0.001 was considered statistically significant, while any test *<* 0.05 at a nominal level of significance was kept for further analysis if post-hoc tests suggested lack of horizontal pleiotropy.

All statistical analyses were conducted in R version 4.00 [47].

### SNP Charaterization and Gene Rhythmicity Analyses

To access rhythmicity, intragenic SNPs (located within a gene) were kept. Each of these genes was input into the cirGRDB database [48]. Rhythmic expression patterns from human and mouse gene expression time-series datasets were queried. Genes analyzed with LSPR [49] and expressed with a period of 22-26 hours, cyclic p-value *<* 005, and amplitude *>* 0.10 were pre-defined as cycling and displayed. Only significant results were kept.

### Gene Set Enrichment

Gene-set enrichment analysis of genes in which protein-coding SNPs reside were subject to gene-set enrichment analyses using the Enrichr platform [**?**]. To address possible phenotype-driven methods of influencing circadian rhythm changes, the gene set was enriched against UK Biobank GWAS and against public GWAS Catalog datasets and visualized in jupyter notebooks; full results are stored: https://maayanlab.cloud/Enrichr/enrich?dataset=54d471c4bb8cf7d38261e595414bc4bd and will be available along with code in our githup repository.

### Reactome Pathway Analysis

CDH11, DENND1A, and FAM76B, along with core clock genes (NR1D1, CLOCK, CRY1, CSNK1D, PER3, PER2, ARNTL2, PER1, TIMELESS, CRY2, NPAS2) were used to form an enriched functional network using Reactome Fi-Viz in Cytoscape. Linker genes (1 degree separation) were included automatically [**?, ?**].

## Results

Mendelian randomization experiments of an exposure to volume change within a specific brain region and an outcome of an evening chronotype resulted in three statistically significant sets of experiments. As shown in figure 1, each region associates with chronotype using varying measures of association. The strongest association is observed in the left anterior Inferior Temporal Gyrus (Z-score = 4.22). This is a positive association, suggesting that an increase in brain volume predisposes to an evening phenotype: in other words, brain volume loss in this region will suggest a morning chronotype. In the left Superior Parietal Lobule, the next strongest association is seen (Z = -2.59), suggesting a brain volume increase in that region associates with a morning chronotype, or a volume loss with evening. The same directionality seen in the right Angular Gyrus (Z = -2.40). Each significant MR experiment was assayed by multiple methods; those with significant results are plotted.

**Fig 1.**
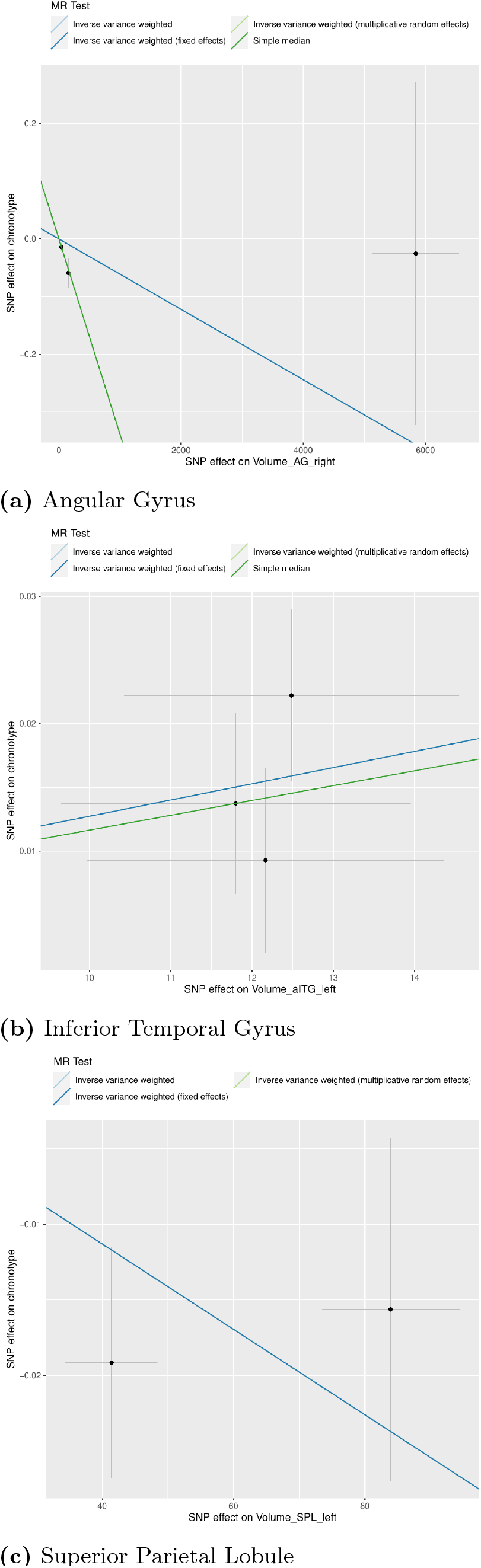
Single nucleotide polymorphisms (SMPs) associating with the Angular Gyrus (AG) - right hemisphere, the left anterior Inferior Temporal Gyrus (ITG), and the left Superior Parietal Lobule (SPL) were used as instrumental variables in 3 Mendelian Randomization (MR) experiments, associating brain volume change with an evening chronotype. Vertical and horizontal lines represent 95% confidence intervals around effect sizes from GWAS studies. AG and SPL volume loss each contribute to an evening circadian phenotype, whereas ITG volume loss associates with a morning preference.

Further details are shown in table 1. Each SNP belonging to significant MR studies has an effect on the exposure (the brain region of interest) larger than the effect on chronotype (panel A), and is annotated to its surrounding gene body if extant, or denoted −if intergenic. The main results of the 3 significant studies are shown in panel B, along with each method used with a *p <* 0.05 threshold score (see methods for significance adjustments). Below (panel C) are heterogeneity and horizontal pleiotropy detection methods, as available. The Egger intercept test (closeness to 0 assumes contradictory pleiotropy) was available for two tests, as three SNPs are required for calculation.

**Table 1.** Mendelian Randomization results:

| roiName | exposure | SNP | Gene | z_exposure | z_chrono |
| --- | --- | --- | --- | --- | --- |
| Superior Parietal Lobule | Volume_SPL_left | rs4836936 | DENND1A | 5.914282 | -2.50240041 |
| Superior Parietal Lobule | Volume_SPL_left | rs7182018 | - | 8.021433 | -1.37700528 |
| Angular gyrus | Volume_AG_right | rs11225471 | RNU7-159P | 8.223570 | -0.08467279 |
| Angular gyrus | Volume_AG_right | rs1532418 | CDH11 | -5.579460 | -2.02652990 |
| Angular gyrus | Volume_AG_right | rs74946541 | - | 5.682191 | -2.32945478 |
| Inferior Temporal Gyrus | Volume_aITG_left | rs12807572 | FAM76B | -5.525641 | 1.27774044 |
| Inferior Temporal Gyrus | Volume_aITG_left | rs62073094 | MAPK8IP1P1 | -6.050975 | 3.29552632 |
| Inferior Temporal Gyrus | Volume_aITG_left | rs79487293 | - | 5.483481 | 1.93747030 |

| exposure | outcome | method | z | pval |
| --- | --- | --- | --- | --- |
| Volume_aITG_left | chronotype | ivw_re | 4.211575 | 2.54e-05 |
| Volume_aITG_left | chronotype | ivw | 3.821073 | 0.000133 |
| Volume_aITG_left | chronotype | ivw_fe | 3.821073 | 0.000133 |
| Volume_SPL_left | chronotype | ivw_fe | -2.589318 | 0.009617 |
| Volume_aITG_left | chronotype | median | 2.433440 | 0.014956 |
| Volume_AG_right | chronotype | median | -2.398811 | 0.016448 |
| Volume_SPL_left | chronotype | ivw | -2.147660 | 0.031741 |
| Volume_SPL_left | chronotype | ivw_re | -2.147660 | 0.031741 |

| exposure | outcome | Q | Q_df | Q_pval | egger_intercept | se | pval |
| --- | --- | --- | --- | --- | --- | --- | --- |
| Volume_AG_right | chronotype | 7.835006 | 2 | 0.0198907 | -0.01618902 | 0.01197142 | 0.4053566 |
| Volume_aITG_left | chronotype | 1.646310 | 2 | 0.4390443 | -0.13307011 | 0.17849368 | 0.5921648 |
| Volume_SPL_left | chronotype | 1.453582 | 1 | 0.2279540 |  |  |  |

(c) Post-hoc tests for horizontal pleiotropy with MR Egger regression intercept test and heterogeneity testing with Cochran's Q suggest little evidence of barriers to direct association, with the exception of AG suggesting possible unseen confounding.

To test if chronotype or sleep duration influenced brain regions under consideration, MR experiments of each trait as exposure vs every brain region under consideration were carried out (table 2). Here, a genetic predisposition to evening chronotype influences the likelihood of increased volumes in the Thalamus and Medial pulvinar nucleus of the thalamus; and a decrease of the superior temporal gyrus (Z = −2.7 − 2.7). Increased sleep duration likewise provides potential causal evidence of increased volume in the central lateral and lateral dosral nucleii of the thalamus (Z = 2.6 −3.9).

**Table 2.** Secondary Analyses: Chronotype and Sleep as Exposures. *ivw*_*r*_*e* and *ivw*_*f*_*e* = inverse variance-weighted average with random- and fixed-effects.

| exposure | outcome | region | method | z | pval |
| --- | --- | --- | --- | --- | --- |
| chronotype | Volume_of_Whole_thalamus_left_hemisphere | Thalamus | median | 2.697455 | 0.00699 |
| chronotype | Volume_aSTG_left | Superior temporal gyrus | ivw_re | -2.653584 | 0.00796 |
| chronotype | Volume_of_PuM_right_hemisphere | Medial pulvinar nucl. thalamus | median | 2.602047 | 0.00927 |
| chronotype | Volume_aSTG_left | Superior temporal gyrus | ivw | -2.425888 | 0.01527 |
| chronotype | Volume_aSTG_left | Superior temporal gyrus | ivw_fe | -2.425888 | 0.01527 |

| outcome | exposure | region | z | pval |
| --- | --- | --- | --- | --- |
| sleep_duration | Volume_of_CL_left_hemisphere | Central lateral nucl. thalamus | 2.710909 | 0.00670991299 |
| sleep_duration | Volume_of_LD_right_hemisphere | Lateral dosal nucl. thalamus | 2.623971 | 0.00869112776 |
| sleep_duration | Volume_of_LD_left_hemisphere | Lateral dosal nucl. thalamus | 3.907241 | 0.00009335598 |

(b) Mendelian Randomization (MR) results with Sleep Duration as exposure and brain regions as outcomes.

Below, we obtained evidence of rhythmicity in several tissues for each target gene. While CDH11 (figure 2 cycles throughout various organs of the body, and DENND1A has been shown to oscillate in stem cells (figure 3, FAM76B cycles not only throught tissues of the body but also in the SCN itself (figure 4a).

**Fig 2.**
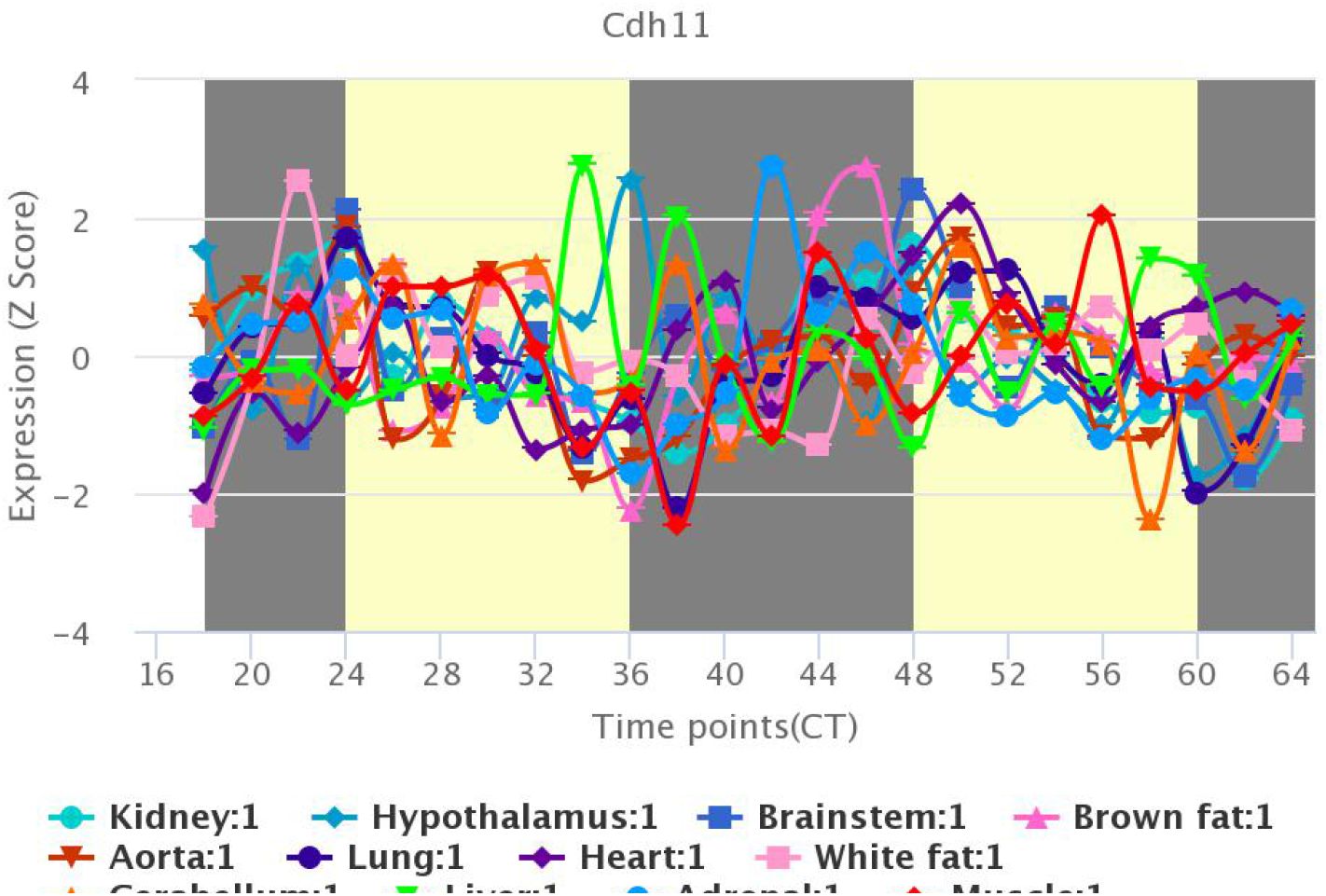
CDH11 exhibits exhibits significant circadian fluctuations across tissues, including the hypothalamus, cerebellum, and brainstem in CirGRDB-drived studies.

**Fig 3.**
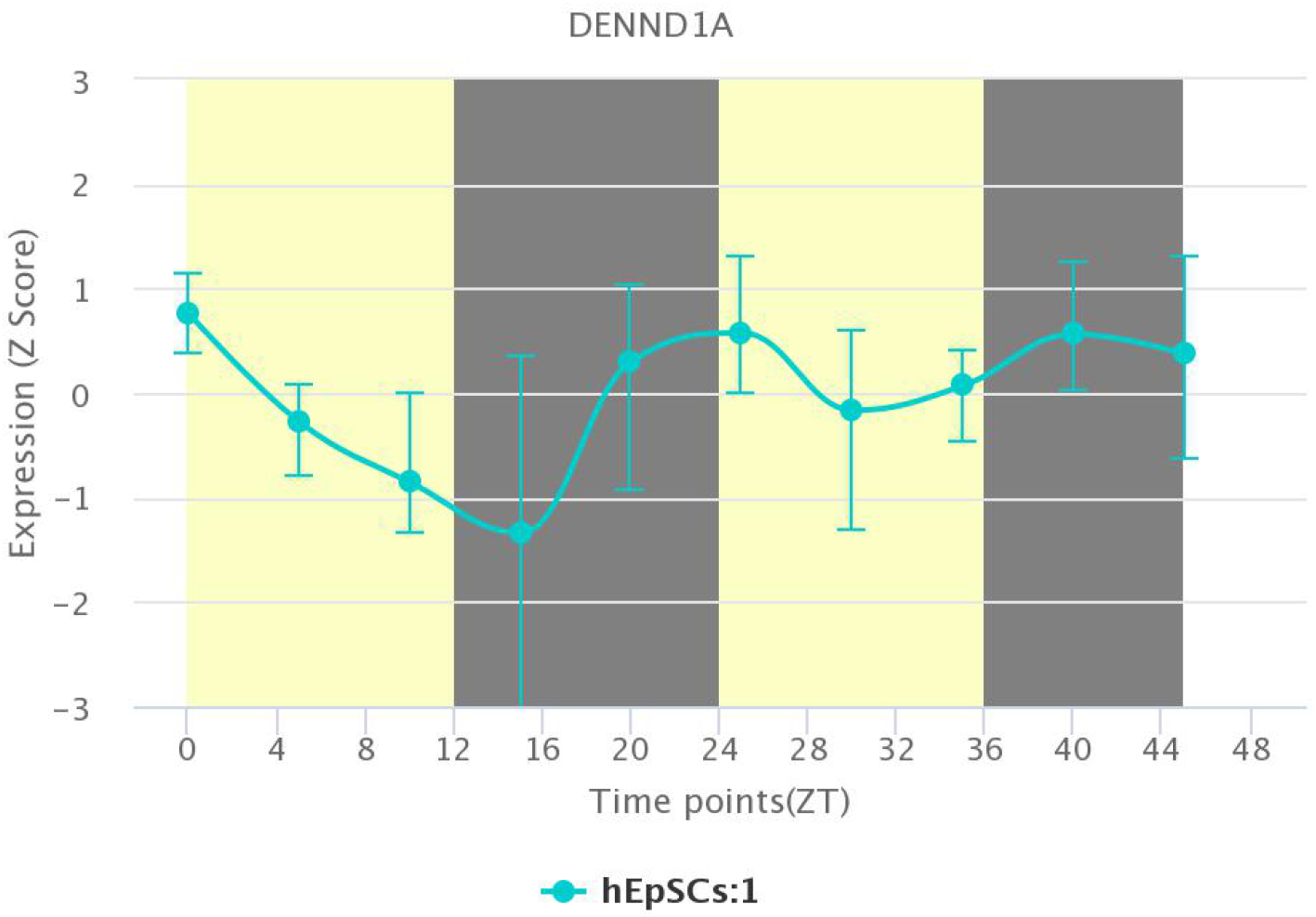
DENND1A shows sinusoidal rhythms in human embryonic stem cells in CirGRDB-drived studies.

**Fig 4.**
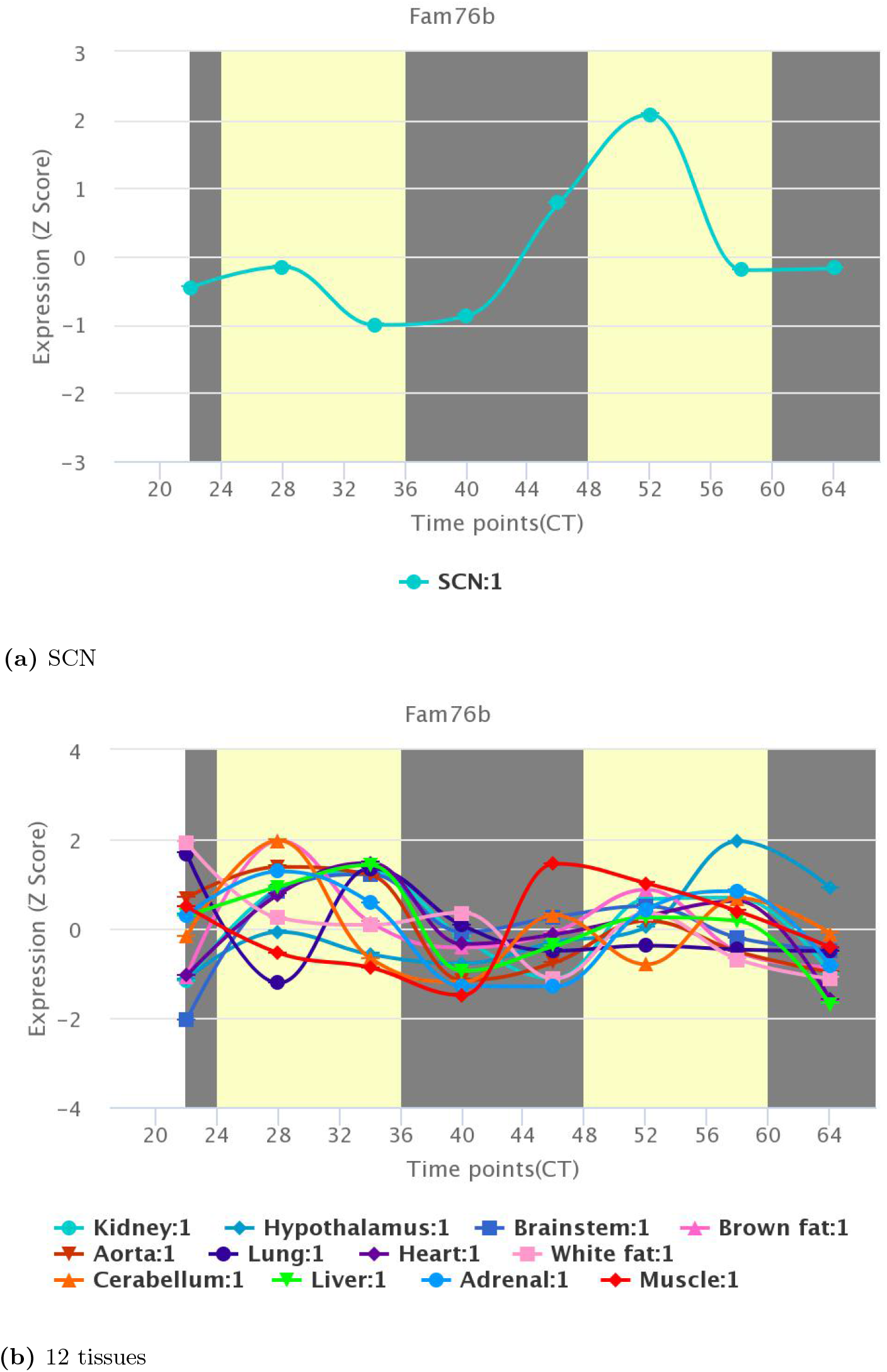
Fam76b oscillates in the SCN specifically and throughout the body generically, indicating specific circadian influenced 24 hour rhythm in several tissues in CirGRDB-drived studies.

**Fig 5.**
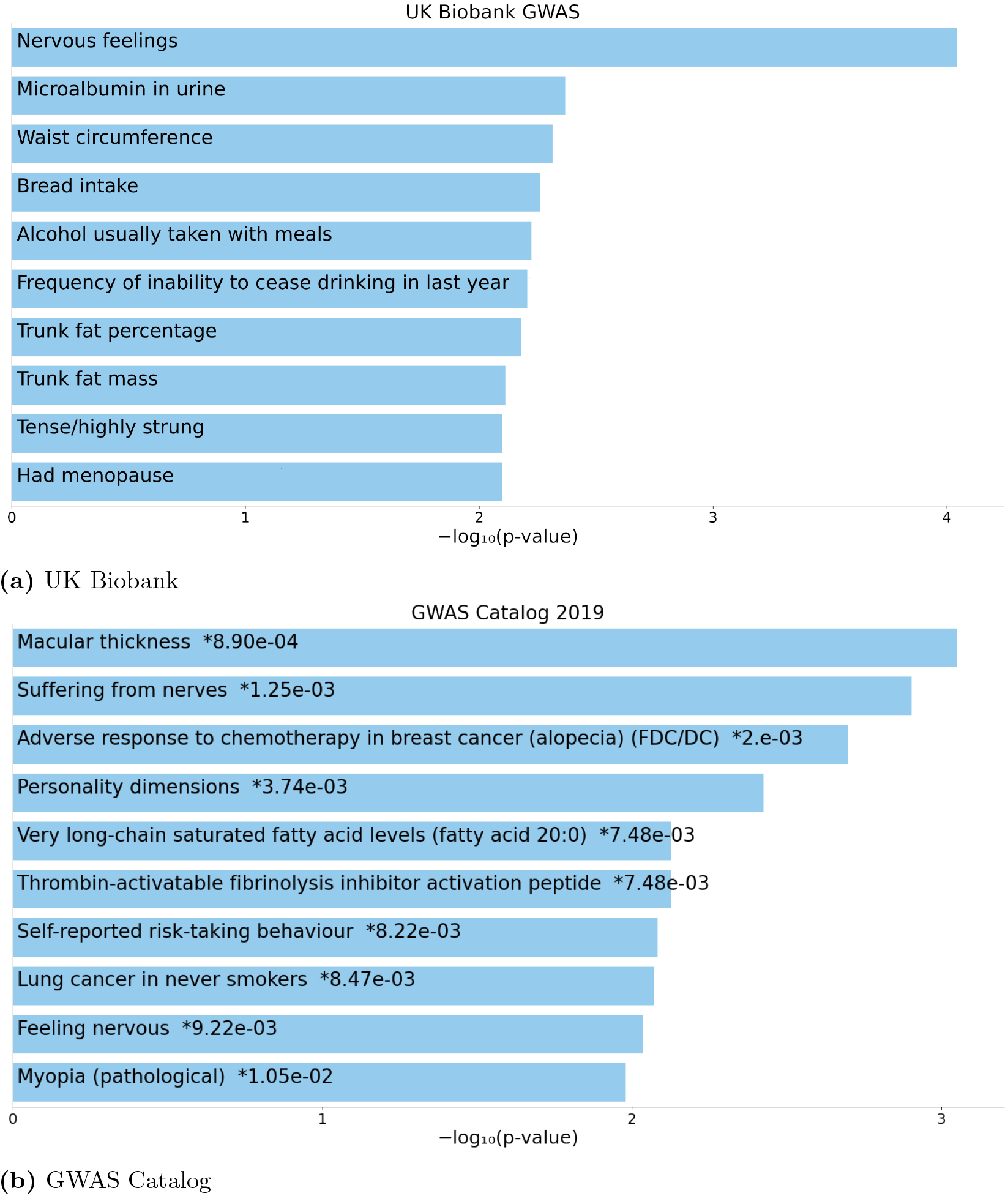
SNPs within zeitgeber genes are over enriched in GWAS studies for chronobiology-influenced traits in the UK Biobank (panel A) and in the GWAS Catalog (panel B).

Each of the three genetic markers was input to enrich existing GWAS study annotations, from UK Biobank and from GWAS Catalog 5. These enrichment methods reveal neuropsychiatric, endocrinological, metabolic, and oncological traits. These traits have independent links to chronotype, and may reveal avenues of influence from these genes’ activities to act as zeitgebers influencing the core clock or circadian phenotype.

Lastly, we projected genes predicted to be non-SCN, intracranial zeitgebers onto Reactome pathways involving cannonical core clock proteins (figure 6. As can be seen, each protein is 1 neighbor away from element of the core clock. FASM76B is connected by an intermediate transcription factor association with ARNTL2 (and thus to the Period proteins). DENND1A is regulated by JUN, which also directionally regulates PER3 and interacts with CTNNB1. CDH11 also interacts with CTNBB1 and EPAS1, a PAS-domain protein that again binds to Period proteins.

**Fig 6.**
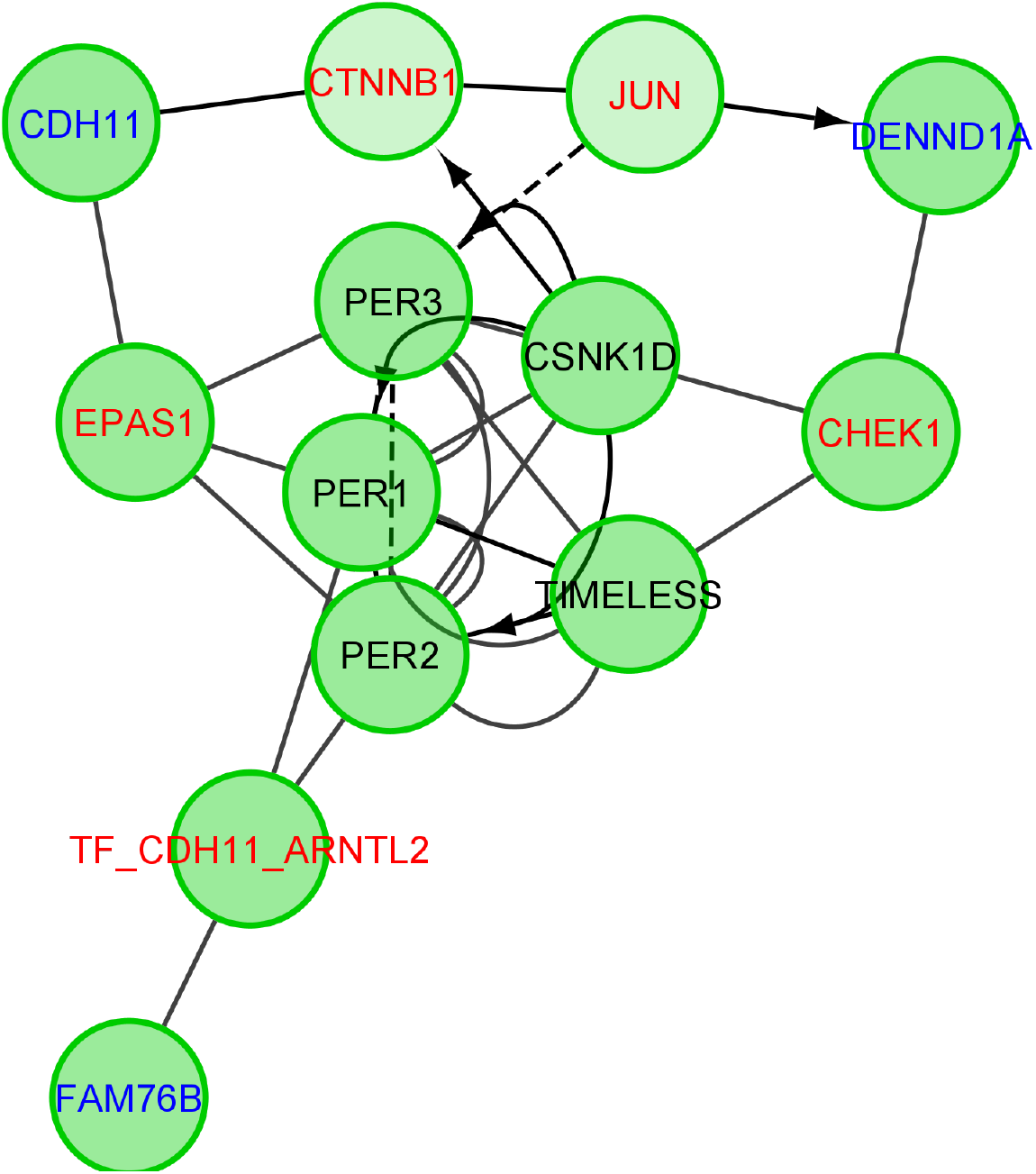
CDH11, DENND1A, and FAM76B all interact on the protein level with secondary messengers of the core clock proteins in the Reactome database. Transcriptional-translational feedback proteins are in black, linking proteins are in red, and external zeitgeber proteins in this study are show in blue.

## Discussion

Using Mendelian randomization and causal inference, the current study examined whether and how genetic variability affecting brain regions outside the SCN might indirectly influence chronotype and potentially relate to some of the chronotype associated phenotypes. We report here three associations with chronotype (morning/evening diurnal preference) outside the SCN based on genetically predicted (FAM76B, DENND1A, CDH11) regional differences in brain volume. Specifically, genetically predicted lower inferior temporal gyrus volume linked to morning phenotype, while lower volume of the superior parietal lobule and angular gyrus linked to evening preference. In addition, evening chronotype exposure influenced superior temporal gyrus volume, and both increased sleep duration and evening chronotype influenced thalamic volume. Thus, our novel findings that genetically mediated associations between measurable brain volumetric regional differences and chronotype exist, suggesting additional (outside the SCN influence the core TTFL) genetic and phenotypic mechanisms, for directly influencing the chronotype. The findings and their implications are systematically discussed below.

### Intracranial zeitgeber genes are rhythmically expressed

Gene/TTFL interactions Possible mechanisms for influencing circadian biology may be endogenous or exogenous, and must take into account molecular, cell, tissue, and organismal views. Our findings suggest evidence of interaction on the molecular level (gene expression/cycling and direct interaction), and from tissue phenotypes under genetic control (Mendelian randomization and phenotypic observations).

As indicated in figures**??**, each gene CDH11, DENND1a, and FAM76B displays some typical sinusoidal rhythms indicative of circadian regulation in non-SCN tissues. DENND1A appears cycling in human-deri1ved stem cells, peaking at 0/24 hours with a trough at 14 hours ZT time - peaking during the beginning of the day (ZT 0) and troughing near the onset of the night (ZT 12) phase during 12 hour light/dark laboratory conditions. No evidence was found in CirGRBD for DENND1A significantly cycling within the SCN. CDH11 and FAM76B appear to have rhythmic expression across several tissues, from the kidney and liver, to brain regions including cerebellum, hypothalamus, and brain stem in the CirGRBD resource. This broad expression of rhythmicity suggests that each of these genes are partially regulated by peripheral clocks which are ultimately regulated by the SCN. A direct mechanism for communication between these clocks, and any two-way relationships, cannot be easily discerned from rhythmic expression in tissues alone. FAM76B, however, also ossilates within the SCN, showing cycling expression in circadian time (CT). Such cycling is observed without reference to external environmental regulators or zeitgebers, indicating endogenous expressed rhythmicity. While such a scale does now allow for easy identification of when (day, night), a double peaking over a 48 hour study is observed (see figure 4.

### Interactions with core clock genes

In figure 6, data from the Reactome database suggests that CDH11, FAM76b, and DENND1A may each be either regulated by, or otherwise interact with, core circadian clock machinery. CDH11 interacts with EPAS1 which itself heterodimerizes with ARNT and interacts with the Period proteins [50]. it also crucially contains the PAS domain along with other proteins core to the central clock - suggestive of common regulation [51]. It also interacts with CTNNB1, which has been shown to bidirectionally interact with core clock proteins [52]. Also known as *β*-catenin, this protein contains an E-box on which the BMAL1-CLOCK complex binds and which CRY/PER complex remove [52]. While the role of the cadherin protein CDH11 within this molecular machinery has not been elucdated, this evidence suggests a potential role. Additionally, a common transcription factor complex regulated by CDH11 and ARNTL2 interacts with FAM76B; this complex likewise interacts with PER2 and 3. Recent invesigations into BMAL1’s regulation of intestinal stem cell signaling show Fam76b significantly cycling in Apc^min^/Bmal1^-/-^ mice when quantification was analyzed by Metacycle [53]. While recently described as a gene of unknown function, Fam76b suppresses inflammation in mice subjected to traumatic brain injury, suggesting a role in both inflammation pathways and repair in neurodegenrative disroders [54]. The protein is expressed in neuronal and glial cell nucleii, and likely resident in macrophages or microglia [54]. While studies of FAM76B in circadian biology are in their infancy, confirmation of presence in neuronal nucleii gives one pilar of necessary support for a role in influencing the central clock itself. DENND1A is a protein associated with polycistic ovary syndrome and possible regulation of androgen synthesis ¡Kempna 2014¿. The gene is expressed in the human cortex ¡cite gtex¿, and members of the connecdenn family of proteins are involved in clathrin-mediated endocytosis [55]. It has been demonstrated that melanopsin (the visual pigment involved in mediating circadian entrainment) colocalizes with clathrin and light-stimulated melanopsis undergoes clathrin-mediated endocytosis [56]. While understudied in a pure circadian context, this may provide an avenue for DENND1A’s influence on circadian biology mediated through the SPL.

### GWAS trait enrichment suggest exogenous influence of chronotype

Intracranial zeitgeber genes were enriched for traits in UK Biobank-exclusive GWAS and in GWAS catalog, suggesting phenotypic roles for these genes which often associate with circadian disorders and chronotype-related phenotypic comorbidity. Among Biobank traits, we note psychiatric (nervous feelings, alcohol misuse, alcohol taken with meals, tense/high strung), metabolic (waist circumference, trunk fat percentage and mass), endocrine (menopause), food-related (bread intake), and kidney health (microalbumin in urine) traits. Several of these traits are associated with chronotype independently of core clock genes. Evening chronotype has a bi-directional (two-way) relationship with alcohol intake [**?**]. Bi-directional MR analyses by those authors show the association is mediated by physical activity, mood, and other external likely zeitgebers. There is a 24-hour rhythm of alcohol craving in adolescent adults, with correlated timings of peak alcohol craving time and an evening phenotype [**?**]. Clock gene expression is altered during the development of drug and alcohol addictions, while circadian disruptions predispose to increased addictive behavior [57]. Beyond this bi-directional relationship, in mice alcohol has been shown to disrupt peripheral skeletal circadian clocks by suppressing core clock genes regulating Bmal1, Period, and Cry genes (orthologs of their human counterparts) [58]. Individuals with generalized anxiety disorder and a morning (early) chronotype reported better sleep quality and parasympathetic emotional regulation than those with an evening preference [59]. There is mixed evidence for supporting a direct association with anxiety and an evening chronotype compared to depression and substance dependence [60], and the directionality of this relationship remains in question. Co-morbidity between substance abuse, including heavy alcohol intake, and mood disorders may introduce mediators of their individual relationships to chronotype [61]; and substance use may serve as a social zeitgeber influencing circadian preference. Obesity and cardiometabolic traits are also show a bi-directional relationship with chronotype; with evening chronotype individuals having a higher body mass index (BMI), ghrelin (hunger hormone) levels, and more unhealthy social eating patterns ¡Erim 2023¿. Meal timing and composition may disrupt endogenous circadian biology and timing of peripheral clocks [62]. Many hormones, including metalonin, cortisol, ghrelin, and leptin have circadian peaks; and while a direct mechanism between feeding time and circadian synchronization is not known, our finding may suggest mediation by the brain regions under consideration [62]. Lastly, there is ongoing debate about the correlation between observed menapause onset age and night shift, which has a strong circadian component [63]. Zeitegeber genes were preferentially over-represented in studies in GWAS catalog. While some echoed evidence from UK Biobank (nervousness, personality, risk taking), others have more explicitly physiological dimensions. We observed associations between these genes and vision-related traits (macular thickness and myopia) and oncology (lung cancer independent of smoking and adverse response to chemotherapy in breast cancer). Like alcohol, links between breast cancer and chronotype may be mediated by cardiometabolic factors already discussed [64]. Taking diurnal preference into account has been suggested as a low-cost adjunctive consideration when planning radiooncology treatments [65]. Regarding lung-specific cancers, again evening phenotypes have an increased predisposition to disease development, partially attributed to social zeitgebers [66]. Pathways from social zeitgeber to direct influence are indirect and as yet unclear. Vision-related traits are clearly mechanistically related to chronotype, since without photoreeptors the primary zeitgeber (light) would be absent. A randomized clinical trail of adults needing cataract surgery demonstrated that melatonin excretion was increased after surgery [67].

Melatonin production, inhibited by light, is secreted from the pineal gland and inhibited by core SCN machinery. In the absence of sight, blind individuals with free-running rhythms can partially entrain their cycles with melatonin administration [68]. Thus, a relationship between macular thickness and melatonin may help explain a circadian influence of macular traits on chronotype. A recent MR study linked a morning chronotype to age-related macular degenration [69]. Elevated melatonin was also obsered in myopic adults, and in a juvenile population myopia was increased with an intermediate or evening chronotype compared to morning [70, 71]. Most of the literature has studied chronotype as an exposure, reflecting the ease of intervention as a therapeutic tool. While these studies did not largely investigate bi-directional relationships between chronotype and traits, the possibility of traits serving as moderators or mediators of brain-region/chronotype influence exists.

An important take-away from these associative observations between chronotype and phenotypes and conditions is that they were derived from our ‘intracranial zeitgeber’ study: from SNPs without a strong chronotype component themselves, embedded in genes which are not independently strongly associated with circadian regulation or entrainment. Potential mechanistic elucidation should take into account not only the snps or gene bodies themselves, but the brain regions that they were associated with.

### Brain regions as independent zeitgebers

Numerous previous studies investigated behavioural and cognitive traits associated with diurnal preference in humans. The published to date studies provide compelling evidence about existing chronotype-dependent differences in cognition (e.g., motor learning, working memory, attention) and link these to time-of day driven differences in cortical excitability and cortical plasticity [72] as well as differences in between and within network functional connectivity [73].

In addition, several recent studies investigated link between chronotype and volumetric brain variations both in adulthood and longitudinally during development. These studies not only found volumetric differences in numerous brain regions, including hippocampus, orbitofrontal cortex, and several subcortical nuclei (nucleus accumbens, caudate, putamen and thalamus) but also strong relationship between some of these volumetric differences to behavioural traits including, loneliness and depression [29–31]. Similarly, Vulser et al. [74] found chronotype driven developmental differences resulting in regional grey matter volumetric differences (mainly in fusiform and prefrontal cortex) resulting in differences in redisposition to self-reported depressive symptoms. It should be noted that all these previous studies specifically examined links between brain/cognitive traits and either self-reported diurnal preference or/and physiological markers of chronotype such as melatonin onset. To our knowledge, this is a first study investigating genetically mediated associations between chronotype and brain regions outside SCN and TTFL.

Each brain region reported here associates independently with chronotype. The three brain regions with volumetric differences linked as implicated by our analysis were: (i) the angular gyrus (AG), (ii) the inferior temporal gurus (ITG) and (iii) the superior parietal lobule (SPL). As shown in figure 1, the strongest signal (and only signal indicating a genetic predisposition to larger volume associating with an evening chronotype) is the left anterior side of the ITG. Using resting-state MRI, Farahani and colleagues investigated whole brain network organization in relation to chronotype by scanning relative time to subject’s waking [75]. Evening preference associated with more small-world type organization. They found particular local changes in the left ITG; interestingly enough these also included some connectivity changes in the thalamus (degree and clustering coefficient) and the left, but not right, AG. They hypothesize that some modes of the visual network have fewer functional paths and connections to other parts of the default mode network 10 hours after waking. The AG was tested by Zhu et al when investigating total sleep deprivation and sustained attention [76]. Slowest reaction times did not peak in near dawn after a night of deprivation, but in around 4am in predawn. The time spent on the test was negatively correlated with activation of the AG, in addition to disruption of the correlation between default node and control networks. While not a direct circadian study, the pre-dawn instead of end-of-study finding does suggest relationships between the connectedness within the structure of the AG and circadian biology. Our findings linking diurnal preference to volumetric differences within the right AG are of particular interests as this area of the brain is not only functionally linked to diverse cognitive processes including attention, spatial cognition, memory, reasoning, and social cognition but has been also implicated as hub interlinking multiple neural networks including the default mode network and other fronto-parietal networks [77, 78],

Decreased connectivity within the default mode network has been previously associated with evening chronotype and implicated as a risk of depressive symptoms [79]. It is plausible that reported here genetically predicted lower volume of the angular gyrus linked to evening preference uderpins the previously reported decreased default mode network connectivity associated with eveningness. Lastly, we found connections between genetic indicators of volume in the superior parietal lobule and chronotype. The Hasher group tested functional activation in older and younger adults, and showed that older adults tested in the morning give better performance and activate the SPL in those tasks [80]. Activation of the left SPL correlated with sleep quality scores when studying sleep deprived nurses who had disrupted circadian rhythms [81]. They observed increased activation of the occipital gyrus on attention tasks in this group. Reske and colleagues [82] performed a small study investigating chronotype and attention to motion tasks; they observed late (evening) chronotype exhibiting attenuation of the SPL activation compared to intermediate but not morning chronotype individuals. Default mode network connectivity was increased in indivuals with restless leg syndrome, a circadian influenced disorder [83]. Working with the same diagnosis group, the stock group showed decreased activation of the SPL by EEG on attention tasks in the evening compared to the morning, hilighting the circadian link to that region’s connectivity [84]. These studies as a whole suggest that connectivity between these regions, and not simply atrophy or brain volume, mediates circadian phenotype and associated traits. In the context of animal studies, there is already strong direct evidence that peripheral oscillators in brain regions outside SCN play a key role in shaping animal behavior and memory processing [12, 85, 86]. While the animal studies specifically explored the influences of the core molecular circadian clock gene oscillation within rodent brain regions, we propose here that parallel can be concluded for genes outside the main TTFL and brain regions implicated by the current study.

### 0.4 Associations with chronotype support direct causation

While the MR results we obtained, upon inspection of directionality of individual instruments, appear to support evidence only of brain morphology influencing chronotype and not the other way around, it was necessary to test if chronotype or related traits also influenced these regions. We performed serial MR experiments of chronotype and sleep duration vs each trait, table 2. Of note, each of these had strong associations with the thalamus or parts thereof; but neither sleep duration nor chronotype GWAS SNPs have likely causal associations with the brain regions we previously identified. This one-way association suggest evidence for causation of some form. Lacking severe heterogenity (showing a uniform directional and statistical signal) further suggest that each genetic signal for all three brain regions is acting in concert - presenting the potential for a molecular or biochemical pathway cascade leading to physiological influence of feedback into clock mechanisms. We tested each analysis for possible horizontal pleiotropy using the MR Egger method, and results do not readily suggest possible colliders, mediators, or moderators in one study, see table 1. Based on the number of SNPs harmonized from the SPL and chronotype, we were not able to conduct plieotropy tests. The Angular Gyrus had the weakest signal, showing results only for a median-based analysis. The presence of horizontal pleiotropy, in this case, would break core assumptions of MR [**?**]. The evidence for homogeneity was weakest in the AG, coinciding with the weakest evidence for causal association. It should also be noted that two genes associated with the Angular Gyrus and Inferior temporal Gyrus respectively by overlapping SNPs in our study were not extensively discussed (RNU7-159P, MAPK8IP1P1); these genes had no functionally annotated products via gene or phenotype ontologies and deserve more time and basic research.

Weaknesses Primary deficiencies in this study include lack of replication GWAS for newly derived brain MRI phenotypes. Ideally, GWAS ‘hits’ will seek replication of at least nominal level to confirm association direction and relative significance. We anticipate such studies being made available to confirm results as more individuals are imaged in the UK Biobank study, and allow us to derive these measurements from their data.

## Conclusion

We have identified three associations with chronotype outside the SCN based on genetically predicted regional changes in brain volume. These associations are between a propensity for brain volume changes leading to an expressed diurnal preference. We have shown that genetically mediated associations between measurable larger regions and chronotype exist, suggesting additional genetic and phenotypic mechanisms for influencing chronotype directly. Reduced grey matter volume in the ITG leads to an increased likelihood of morning phenotype, while reduced volume of the SPL and AG lead to evening preference. Evening chronotype exposure influences changes in the STG, and both increased sleep duration and evening chronotype preference influence thalamic nucleii volume. Phenotypic and genetic interaction evidence points to potential zeitgeber mechanisms, specifically around FAM76B, DENND1, and CDH11 and their potential interactions with the core clock transcriptional-translational feedback loop.

## Supporting information

Supplemental information, including full GWAS results and lists of all regions studied, will be included in future versions.

## Acknowledgments

This research has been conducted using the UK Biobank Resource under Application Numbers 29447 and 31224. The computations described in this paper were performed using the University of Birmingham’s BlueBEAR HPC service, which provides a High Performance Computing service to the University’s research community. See http://www.birmingham.ac.uk/bear for more details.

## Disclosures

John A. Williams is currently an employee of Eisai, Inc. Eisai, Inc had no role in funding or design of this study.

References

yes

